# Electrical Stimulation Rejuvenates Tunicates: Altered Stem Cell and Immune Activity

**DOI:** 10.1101/2025.05.17.654683

**Authors:** Jos Domen, Yotam Voskoboynik, Tom Levy, Erica M. Domen, Katherine J. Ishizuka, Karla J. Palmeri, Chiara Anselmi, Thomas Rolander, Norma F. Neff, Angela M. Detweiler, Irving L Weissman, Kimberly L. Gandy, Debashis Sahoo, Ayelet Voskoboynik

**Author notes:** Equal senior contribution (Jos Domen, Kimberly L. Gandy, Debashis Sahoo, Ayelet Voskoboynik). Corresponding Author E-mail: Ayelet Voskoboynik Jos Domen Debashis Sahoo.

## Abstract

Applicable methods of rejuvenating organisms and improving resistance to environmental stimuli are needed. During attempts to synchronize heart rates in unhealthy colonial chordates, we observed morphological rejuvenation. While the importance of endogenously generated bioelectric currents in development is well-established^1,2^, and exogenously applied current has shown promise in regenerative medicine^3,4,5,6,7,8,9^, a model that robustly increases longevity and fertility while providing detailed mechanistic insights has not been reported. Here, we report the establishment of such a model using pulsatile electrical current (PEC) in *Botryllus schlosseri*, an established colonial chordate model^10,11,12,13,14,15^. PEC treatment significantly improved survival, morphological integrity, stem cell mediated regeneration, and gonad production in *Botryllus*. Transcriptomic analysis revealed pathway changes associated with cellular metabolism, cell cycle, stem cell activity, DNA repair, and immune modulation. Notably, PEC-induced expression patterns resemble the exercise-induced macrophage-associated transcriptional response previously observed across several mammalian species^16,17^. This transcriptomic signature correlated with an increase in immune-cell-containing populations. These findings demonstrate that PEC can improve longevity, vitality, and reproduction in an established model renowned for defining broadly applicable biological principles. These studies offer insights into novel strategies for promoting healthy aging and organismal survival.

## Introduction

Studies that lend insight into improving fertility, longevity, and resistance to changing environmental conditions are critical. We have designed a system to evaluate these conditions in *Botryllus schlosseri,* an organism whose asexual life cycle allows high-throughput evaluation. *B. schlosseri* is a member of the subphylum Tunicata (Urochordata), the closest living sister group to Vertebrata (within the phylum Chordata)^12,18^. The *B. schlosseri* genome exhibits significant sequence similarity to ≈ 75% of human protein-encoding genes^12^. As such, *B. schlosseri* serves as a model for pre-vertebrate evolution and a translational model for a broad range of organisms. Indeed, *B. schlosseri* has been successfully used to study stem cell aging and stem cell competition^14,19,10,15,20,21,22,27,32,35^, with findings that have led to discoveries in their mammalian counterparts. Our new model evaluating pulsatile electrical current (PEC) in *B. schlosseri* may have profound implications for fertility and longevity and the study of responses and resistance to environmental challenges in a wide range of organisms.

*B. schlosseri* consists of one or more systems (“flowers”), which are composed of zooids (“petals”) and buds (Fig.1a-d). The entire colony is enclosed in a gelatinous tunic. Each zooid has a heart that provides the pulsatile force to support circulation in the shared vasculature of a colony. The vasculature ends in ampullae, which are visible in the periphery of the tunic. *B. schlosseri* has well-characterized sexual and asexual reproductive cycles^23^. In the asexual reproductive cycle, genetically identical zooids mature from buds originating from the predecessor zooid^23,24,25^.

Every week, primary buds produce one or more secondary buds. After one week, these secondary buds mature into primary buds, which simultaneously develop into adult zooids. The older zooids undergo apoptosis and resorption, being replaced by the newly emerging adult zooids in a “takeover” process. This once-per-week asexual reproductive cycle (20^0^C) allows *B. schlosseri* to be maintained and expanded for decades in the lab on microscope slides, making the organisms ideal for systematically evaluating factors that alter biological properties^26^. Indeed, our oldest colonies have been maintained in the lab for more than 24 years, having gone through takeover more than 1,000 times. While it is not clear how long individual colonies survive in the ocean, the availability of this age range of colonies in our lab allows us to compare young animals with those cultured for decades (Fig.1a-d). During the March through October reproductive season (Monterey Bay, CA), when colonies are sexually mature, zooids develop gonads that produce oocytes and/or sperm and can initiate the sexual reproductive cycle. This cycle, however, happens with relatively low frequency in culture compared to the wild. In the sexual reproductive cycle, fertilized eggs within the zooids undergo classic chordate development, producing tadpole-shaped larvae. These larvae disperse, settle on a substrate, undergo metamorphosis (which includes tail and partial nervous system loss), and become sessile filter feeders^23^.

As colonies age, both sexual and asexual reproduction slows, demonstrating reduced regenerative potential^10^. Correlative morphological changes associated with aging include increased pigmentation, altered blood vessel shape, and significantly decreased zooid size (Fig.1a-d). Significant molecular changes accompany these morphological changes, including changes in the expression patterns of genes associated with circadian regulation^21^. Additionally, neuron numbers within the *B. schlosseri* central nervous system (CNS) decrease with age, and these neurons exhibit significant changes in gene expression linked to human neural stem cells and neurodegeneration pathways^27^.

Previously, a system was tested for providing pulsatile electrical current (PEC) to cardiomyocytes derived from mouse embryonic stem cells via a clinical-grade pacing device (Medtronic 5375). Cardiomyocyte contractions increased after pulsing at higher rates with the Medtronic pulse generator, confirming the power of this approach (SI Fig.1b). We adapted this system for use in *B. schlosseri*, hypothesizing that increased or coordinated heart contractions and the resultant increased circulation may alter biological properties. Early experiments demonstrated striking morphological improvements following PEC exposure in young and old colonies, prompting further exploration of PEC in aging.

Naturally occurring bioelectric currents play a critical role in development and regeneration. Studies have revealed that all cells generate electrical networks that can control gene expression and behavior. ^1,2,28,29^. In invertebrates such as *Planaria*, endogenous bioelectric currents are critical for regeneration, while in amphibians, like salamanders, such currents are crucial for tail regeneration. These principles extend to mammals, where several applications are being studied. For instance, electrical modulation of neural stem cells is being evaluated to improve function^3^, and electrical stimulation is used to treat peripheral nerve and spinal cord injuries^4,5^. Clinically, externally generated currents have shown promise in wound healing^7^, immunomodulation^8,9^ and the regeneration and repair of nerve, bone, cartilage, and muscle^6^.

Decades of fieldwork by investigators worldwide have explored the use of electrical current to promote regeneration in marine corals, yielding promising results (reviewed in^30^). However, studies of the associated mechanisms are lacking^31^. Despite the vast number of studies on bioelectrical stimulation, limited data exist on the direct application of electrical current to alter organismal stem cell-mediated regenerative activity, longevity and fertility. To our knowledge, this is the first study evaluating a reproducible, broadly applicable PEC regimen that can be studied with scientific rigor in the context of stem cell activity, longevity and fertility. Our research extends beyond observational studies to include detailed cellular and molecular responses.

Data in humans has shown that myeloid populations have transcriptomic shifts with stress or exercise. Exercise induces an immediate surge in macrophage-associated transcriptomics consistent with a pro-immune response M1 activation, with an equal and opposite long-term balancing effect consistent with the cell repair focus M2 activation. This finding was consistent across different species (mouse, rat, and human), tissues and sampling methods^17^. Our model system abruptly increases the heart rate of *B. schlosseri* above its normal rate of 60-90 bpm^21^, consistent with what organisms may experience with stress or exercise. We documented correlative transcriptomic changes in our PEC-*Botryllus* model.

Our *B. schlosseri* studies establish that PEC enhances morphology, regeneration, and lifespan. Furthermore, PEC expands cell populations that contain immune cells and induces transcriptomic changes that co-opt stem cell and immune activity to enhance reproduction. This work will provide critical data and a model system to unify efforts from different research fields, which is essential for defining broadly applicable methods to address critical and time-sensitive challenges in our world.

## Results

### PEC induced younger phenotypes: Initial observations and regimen development

Initial observations suggested that PEC treatment caused unhealthy or older animals to assume a younger phenotype, noticeably within 48 hrs. A few weeks after pulsation, these animals exhibited features characteristic of younger individuals, including brighter pigmentation, more elongated ampullae extending to the tunic edge, a clearer tunic, and increased gonad production (Fig.1e-f). For PEC application, we used a Medtronic 5375 pulse generator, consistent with devices used in clinical cardiac surgery (SI Fig.1a). Our PEC regimen was developed with a strong foundation in clinical practices, specifically leveraging existing knowledge of safe and effective voltage and frequency parameters employed in cardiac pacing. To evaluate various PEC regimens for *B. schlosseri*, we pulsed animals at 20 mA and 150 pulse per minute (ppm)— a rate faster than the typical *B. schlosseri* heart rate of 60 to 90 beats per minute (bpm)^21^ — for duration ranging from 5 minutes to 16 hours. After evaluating pilot experiments (SI Table 1), a regimen with 20 mA, 150 ppm for 5 minutes, repeated 3 times with 20-minute intervals was chosen for subsequent studies. For some initial experiments, 20 mA, 150 ppm for 5 minutes was applied 6 times with 48-hour intervals. In our experiments, young colonies ranged from 2 to 18 months, while old colonies had been maintained in mariculture for 14 to 24 years.

**Fig 1.**
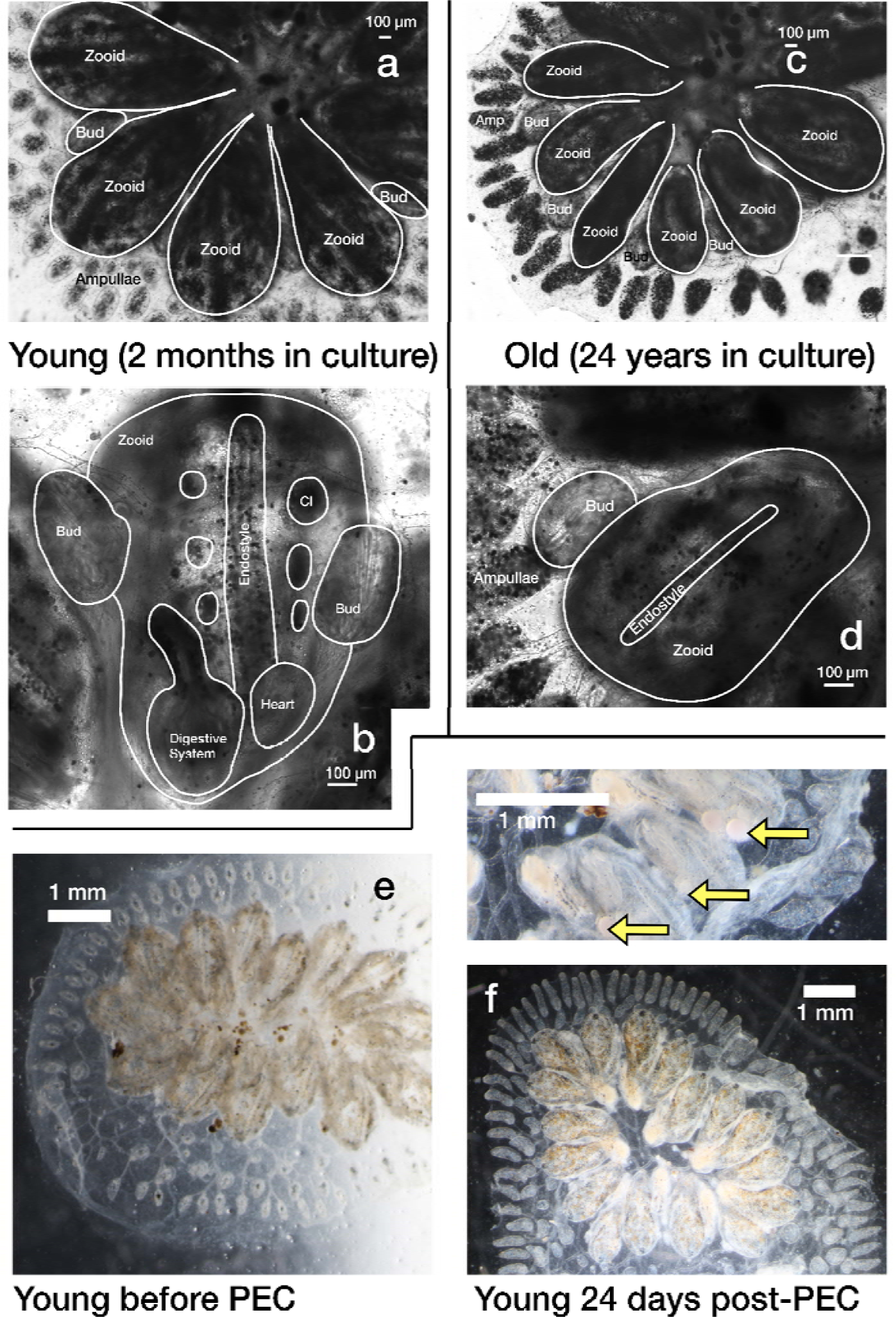
Age-related morphological changes and PEC-induced rejuvenation in *Botryllu schlosseri.* (a-b) Live images of a young (2-month-old) colony. (c-d) Live images of an old (24-year-old) colony. (a) A *Botryllus* colony comprises interconnected zooids, each bearing buds and connected to a shared blood vessels network terminating in peripheral ampullae (Amp). Young colonies (a-b), exhibit larger sized zooids with less pigmentation compared to old colonies (c-d), which also show differences in ampullae’ morphology, size, and pigmentation^21^. (b) A young zooid, requiring two 10x magnification images to capture its entirety. Internal organs such as the digestive system, heart, endostyle, and cell islands (CI) can be observed through its semi-transparent body. (d) An entire old zooid and its buds at the same developmental stage and magnification fit within a single frame, highlighting the significant size reduction in older zooids and their organs. (e) A 10-week-old young colony before exposure to PEC, shows an opaque tunic, retracted ampullae, and the absence of gonads and embryos. (f) The same colony 24 days after PEC exposure shows a transparent tunic, extended ampullae, and embryos and gonads in nearly every zooid (arrows). Scale bar = 100μ*m* (a-d) or 1mm (e-f). Magnification: a, c: x4; others: x10. Zooids, buds, and internal organs are outlined.

### PEC increased sexual and asexual reproductive

We had appreciated that pulsatile current had positive consequences in some organisms, but we did not know if this correlated with heart rate changes. Our findings show that PEC-treated colonies (3×5min) exhibited an increased heart rate 1-2 hours following stimulation, which persisted, albeit to a lesser extent, up to 18 days after treatment (Fig. 2a and SI video 1). Furthermore, PEC treatment enhanced reproductive output in young colonies. Specifically, PEC-treated young colonies produced significantly more zooids than controls by day 23 (Fig. 2b). Notably, gonads were observed in 7 out of 9 treated colonies compared to only 1 out of 9 untreated controls, *p*=0.0115 (Fisher exact test) (Fig.2c).

**Fig 2.**
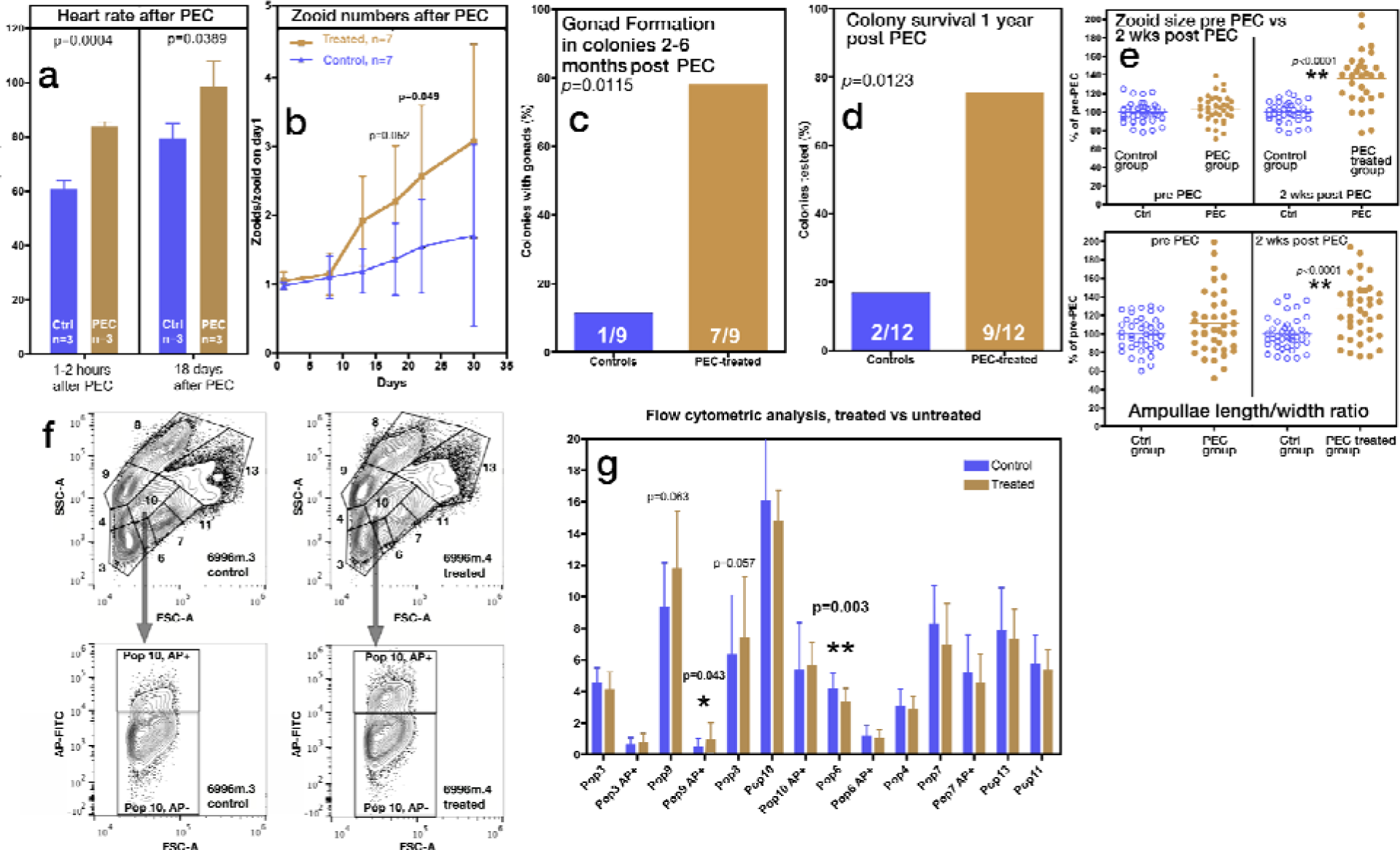
PEC treatment enhances physiological function and cellular composition in young colonies. (a). Comparison of heartbeat frequency between control and PEC-treated colonies (3×5min) on the day of treatment (Left) and 18 days later. Data are presented as average ± SD (unpaired t-test). (b) Average zooid numbers over 35 days for 7 untreated control colonies and 7 PEC-treated colonies (4 to 5 months old at treatment). Zooid numbers in treated colonies was significantly higher on days 20 and 25 following treatment (unpaired, t-test). (c). Proportion of zooids with gonads in treated and controlled animals 2 to 4 months post-treatment. A larger proportion of treated animals developed gonads in most of their zooids. (d). Survival rates of 12 control and 12 treated colonies one year after treatment. (e). Average zooid area and ampulla length/width ratio measured using ImageJ two weeks after treatment. Data are shown from 4 colonies per group, with 10 measurements per colony. Values are normalized to the untreated control group ** p<0.0001. (f) Flow cytometric evaluation: Gating setup according to^32^. Gates are based on scatter parameters, propidium iodide staining (dead cell exclusion), and a fluorescent substrate for alkaline phosphatase (AP-FITC) (g) Comparison of blood cell populations between control (blue bars) and PEC-treated animals (twine bars) 4 weeks post treatment. Significant differences were observed in population 6 and population 9 AP+ (contains Morula cytotoxic cells). Borderline non-significant differences are also indicated. Figure 2f shows an example of the ability to separate population 10 based on AP^+^ and AP^-^ status.

### PEC extended lifespan and rejuvenated health indicators

PEC-treated colonies thrived long after the treatment ended and showed increased lifespans. At 12 months post-stimulation, 9 out of 12 stimulated colonies were still alive, undergoing weekly budding cycles, whereas only 2 out of 12 control colonies survived that long (*p*=0.0123, Fisher’s Exact Test) (Fig.2d). While difficult to quantify precisely, PEC also resulted in more robust blood flow within colonies and reduced hyperpigmentation, especially in the ampullae (SI video 1). We also observed that the size and length/width ratio of zooids and ampullae were significantly larger when comparing PEC-treated colonies with control colonies (p<0.0001) (Fig.2e), consistent with phenotypes seen in younger animals.

### PEC rejuvenates aged *Botryllus* colonies

PEC applications improved the morphology and circulation in older organisms (14 and 24 years old), reducing hyperpigmentation and increasing zooid numbers closer to levels closer to those typically found in younger organisms (Fig.3). For example, Fig.3a-d shows these results in a 24-year-old colony (944axBYd196-6-4), while panels 3e-h show similar data for a 14-year-old colony (SC109e). It’s noteworthy that these older colonies typically regenerate smaller zooids compared to young colonies (as demonstrated in Fig.1a-d; using untreated 944axBYd196-6-4). However, following PEC treatment, the size of these older zooids significantly increases (Fig.3d and 3g).

**Fig 3.**
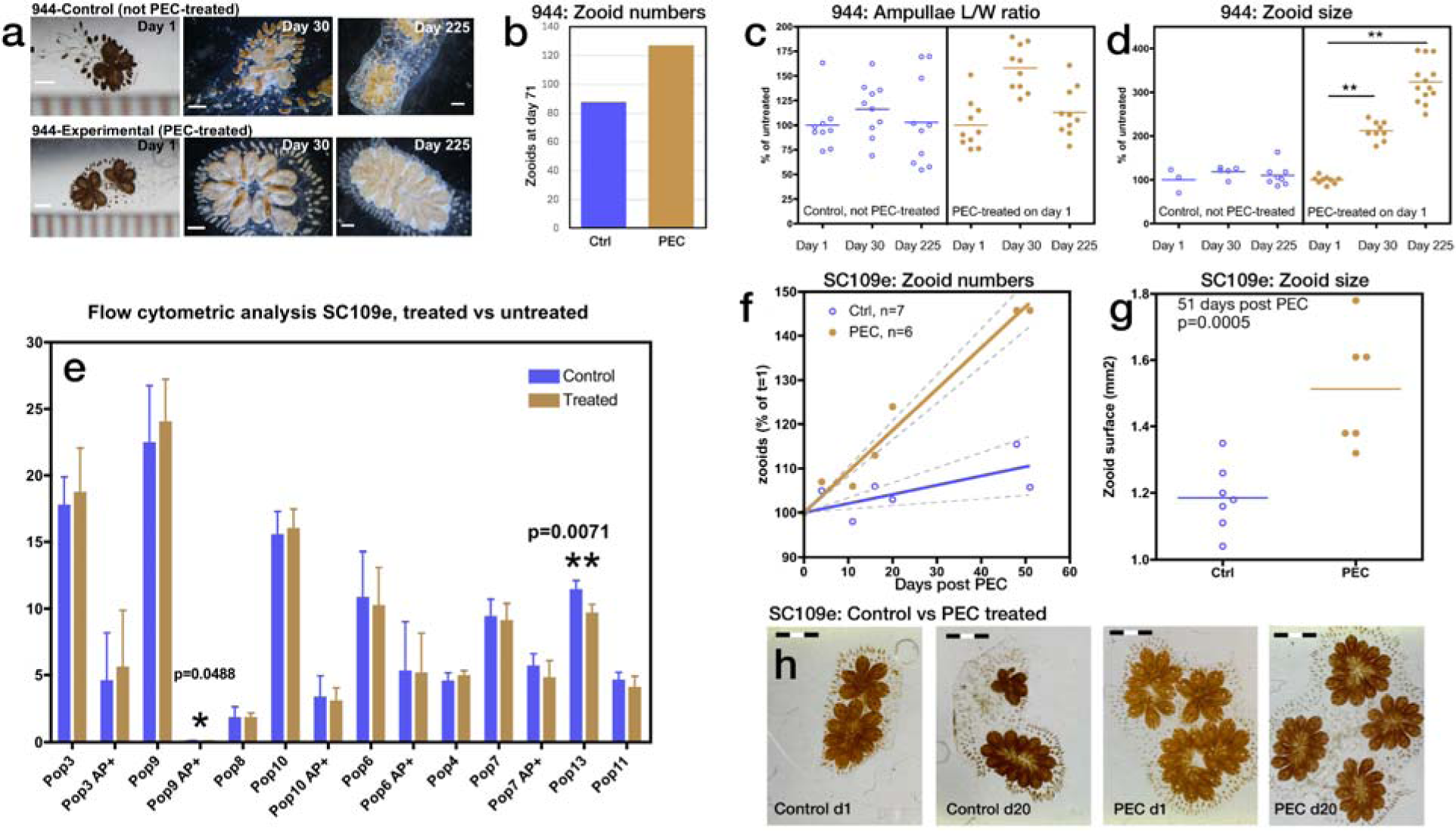
PEC treatment rejuvenates aged *Botryllus* colonies. (a-d) Effects of PEC treatment on colony 944, 20+ years old. (a) Prior to PEC treatment this colony exhibited poor condition (criteria as in Fig. 1). A single PEC treatment resulted in clear short-term and long-term improvement. (b) Zooid number over time in a control and PEC-treated colony. Data suggested an increased generation of zooids in the PEC-treated animals (data from a single colony per condition) (c) The length/width ratio for the ampullae does not change significantly. (d) Zooid sizes increased significantly after PEC treatment (**: p<0.0001). (e-h) Effects of PEC on SC109e (14 years old). (e) Flow cytometry analysis (gating as in Fig.2f) of SC109e, 7 weeks post-treatment. Similar to young colonies (Fig.2), PEC-treated animals show significantly more AP+ cells in population 9 (which contains Morula cytotoxic cells) and population 13 (which contains phagocytic, macrophage-like cells). n=3 (control) and n=2 (treated) animals per group. Panels (f) and (g) Zooid number (f) and size (g) in control and treated SC109e colony. Dashed lines indicate 95% confidence intervals. (f) PEC treatment at t=0 (h) Morphology of clone SC109e on days 1 and 20 after PEC treatment. Scale bars are in mm.

### PEC induces dynamic shifts in *Botryllus* immune cell populations

To investigate the effects of PEC on *Botryllus* cell populations we employed flow cytometry, building on previously characterized stem-cell and immune cell-containing populations in *B. schlosseri*^32^ which were identified using propidium iodide (PI), size, granularity, and a FITC-labeled substrate for Alkaline Phosphatase, an enzyme expressed at high levels in some stem cells^32^. The gating setup for *B. schlosseri* cell populations^32^ in a control colony and a PEC-stimulated colony is shown in Fig. 2f. Our analyses revealed a significant increase in populations of immune cells, including phagocytes and cytotoxic cells, two months after PEC stimulation. Notably, both young and old animals have significantly higher numbers of AP^+^ cells in population 9 at 14-33 days post PEC (Fig.2g; p < 0.05). This population encompasses morula cells, which are implicated in cytotoxic activity within *B. schlosseri* colonies^32,33^. Population 6, which contains lymphocyte-like cells^32^, is significantly decreased in PEC-treated young colonies. A statistically significant decrease is also observed in population 13 in aged colonies, which comprises phagocytic cells (Fig.3e). Two other populations, including population 9, show increased numbers which reach borderline significance in young colonies (p = 0.057-0.063), as indicated in Fig.2f. It is important to note that flow cytometric analysis in *B. schlosseri* is limited by both a lack of antibodies and high levels of background fluorescence, but our long-term experience in studying of *B. schlosseri* cell populations has allowed us to use the existing tools effectively for this analysis.

### Transcriptomic reprogramming by PEC: lasting effects on metabolism and stem cell pathways

PEC triggers long-lasting gene expression changes, affecting cellular metabolism, cell cycle, stem cell and immune activity. In old *B. schlosseri* colonies, among the transcripts differentially upregulated 24 hours after PEC we identified genes associated with the circadian clock, transcriptional activation of mitochondrial biogenesis, metabolism of steroids, and telomere maintenance pathways (Fig.4b; SI Table 2). Notably, at 50- and 90-days post-pulse, these older colonies show differential upregulation of key signaling pathways, including Wnt, Slits/Robo, and VEGF (Fig.4b; SI Table 2). Distinct transcriptomic responses were observed in young colonies. Twenty-four hours post PEC treatment, upregulated transcripts were associated with metabolism, the complement cascade and regulation of insulin-like growth factor. Following this, TGF-beta receptor signaling and the Notch pathways showed upregulation at 13- and 50-days post-PEC. One hundred thirty-two days after treatment, pathways involved in protein metabolism, EGFR, IL-17 and innate immune system signaling pathways are upregulated in these young colonies (Fig.4a; SI Table 2). These changes align with the observed long-lasting improvements in morphological and cellular characteristics, as well as the enhanced stem cell mediated regeneration activity observed following PEC treatment.

**Fig 4.**
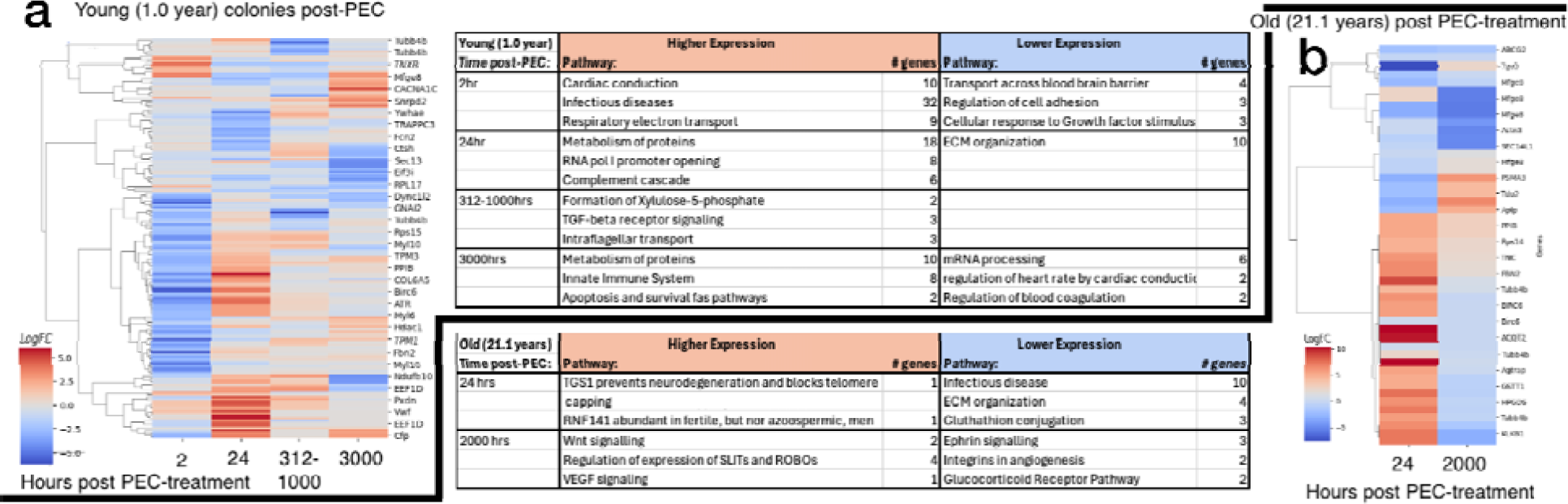
Temporal gene expression changes following PEC treatment. Relative gene expression changes (log fold change) between PEC-treated and untreated colonies at various time points. (a) Young colonies: 2 hours, 24 hours and >13 days after PEC. (b) Old colonies: 24 hours and 83 days after PEC. Differentially regulated genes are associated with diverse cellular processes, including circadian clock, mitochondrial biogenesis, steroid metabolism, protein auto-ubiquitination, Wnt signaling, Slit-Robo signaling, and VEGF signaling. Complete lists of differentially expressed genes are available in SI Table 2.

### PEC induces a conserved M1-to M2 macrophage like transcriptomic shift in *Botryllus*

The effects of PEC treatment in *B. schlosseri* mimic an exercise-induced macrophage activation transcriptome signature observed in mammals. In previously reported mammalian studies (humans, rats, and mice), exercise induces a dynamic shift in the macrophage activation transcriptome signature. Violin plots in Fig. 5 display this transition, representing the analysis of 338-gene sets that identified macrophage polarization states, as previously defined^16^. This shift involves a transition from an initial pro-inflammatory (M1) state observed within 6 hours post-exercise in mammals (Fig. 5a), to a subsequent reparative (M2) state that favors tissue repair, occurring 48 hours and beyond post-exercise^17^ (Fig. 5b). In our *B. schlosseri* model, 112 of the 338 macrophage polarization state genes had identifiable homologous (SI Table 2). Comparative analysis reveals that 2 hrs after PEC stimulation, *B. schlosseri* exhibited an initial M1-like transcriptome signature Fig. 5c (column 1) similar to the early response seen in humans after exercise (Fig. 5a). Subsequently, *B. schlosseri* transitions to an M2-like phenotype by approximately 300 hrs post PEC treatment (Fig. 5c). This transition to an M2 state, which mirrors that observed in human after consistent exercise (Fig. 5b), occured in both young (Fig. 5c, columns 2 and 3) and old colonies (Fig. 5c, column 4). For both age groups, this M2 phenotype persisted from 300-1000 hrs (∼41 Days) following PEC treatment, trending back towards baseline after 42 Days.

**Fig 5.**
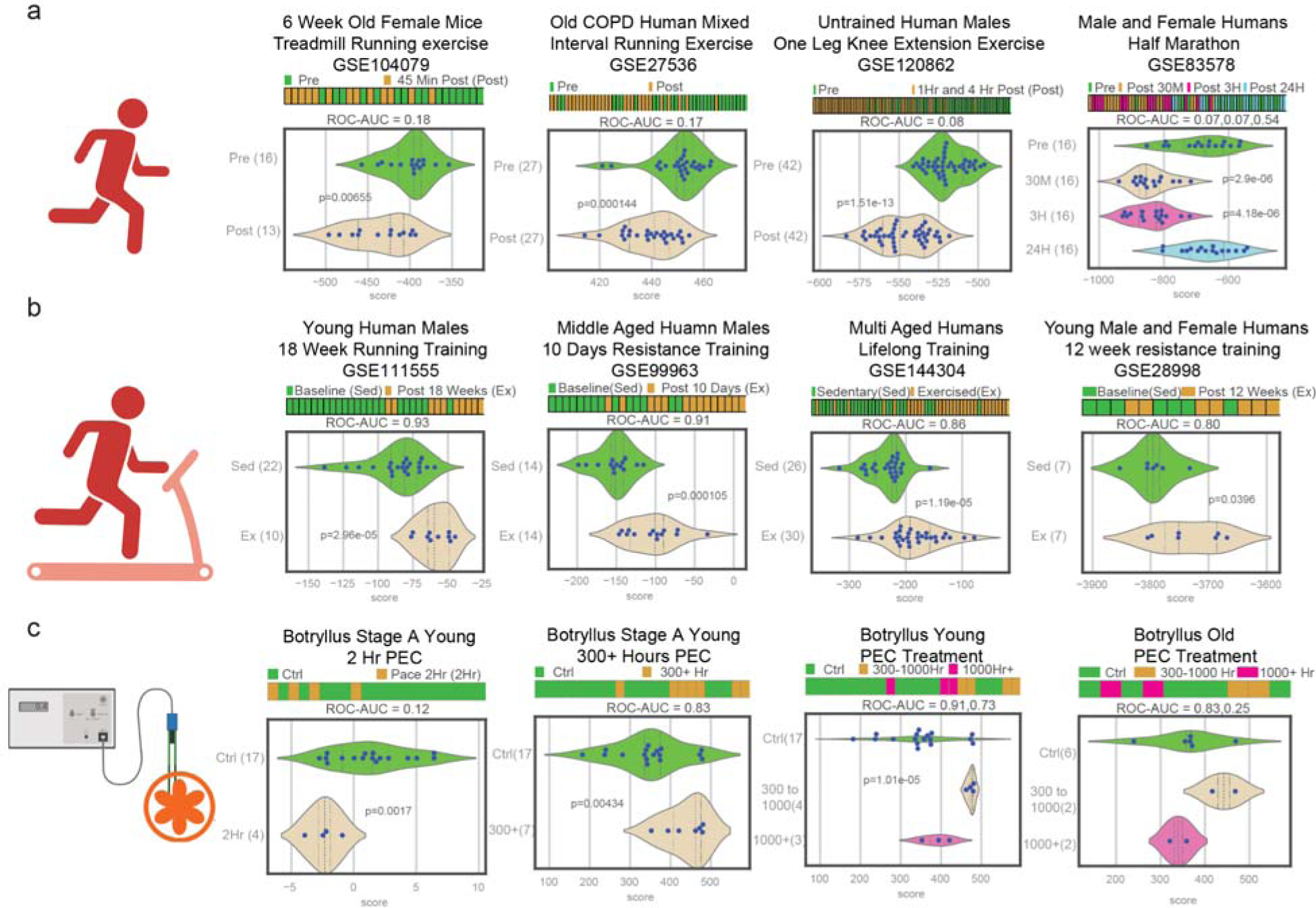
Macrophage polarization associated signature in exercise and PEC. (a-c) Macrophage transcriptional profiles were analyzed using a 338-gene model (M1: 48 genes; M2: 290 genes^16,17^, with 13 M1 and 133 M2 *Botryllus* homologs. M1-associated genes were assigned negative weights, and M2-associated genes were assigned positive weights. Each sample was assigned a score based on the sum of the average z-score multiplied by the weight of all the model genes, allowing for comparison relative to controls. Thus, higher M1 gene expression resulted in lower scores, while higher M2 gene expression resulted in higher scores. (a) In humans and mice, 6 hours post-exercise, scores were significantly lower compared to unexercised controls, indicating an immediate M1 phenotype. (b) In humans, at least 48 hours post-exercise, scores were significantly higher than unexercised controls, indicating a sustained M2 phenotype. (c) *Botryllus* exposed to PEC exhibited similar macrophage polarization patterns. In young colonies, 2 hours post-PEC, scores were significantly lower than controls, indicative of an M1 phenotype. At least 300 hours post-PEC in all young samples, scores were significantly higher compared to controls, indicative of an M2 phenotype. The strongest M2 signal was observed between 300- and 1000-hours post-PEC, declining to control levels thereafter. This pattern was observed in both young and old colonies, although statistically significant only in young colonies (likely due to larger sample size). The y-axis displays the experimental groups, including the number of samples within each group. P-values are shown for significant samples. Complete M1 and M2 Botryllus-associated gene lists are available in Supplementary Table 2. a and b adopted from (17).

## Discussion

The dramatic and persistent rejuvenating effect of PEC in aged *B. schlosseri* colonies strongly suggests a fundamental impact on stem cells, as organismal aging in this species is driven by the aging of its stem cells. Using lineage tracing and transplantation, we have previously isolated stem cells, identified stem cell niches, and demonstrated the migration of both germline and somatic stem cells from adult zooids into developing buds^15,20,35,32,23^. These studies established the crucial role of stem cells in *B. schlosseri* bud development. During the weekly replacement of old zooids by new generations, stem cells are the only cell population that persists throughout the colony’s lifespan. This unique characteristic allows for the study of stem cell aging by observing the phenotypes of tissues and organs that develop de novo each week from these long-lived stem cells.

Stem cell aging is also a recognized feature in mammalian systems; for instance, hematopoietic stem cells (HSCs) expand with age but exhibit a myeloid bias (my-HSC)^36^, and depletion of these my-HSC can restore a more youthful immune system^37^. While *B. schlosseri* currently lacks the specific purification markers available for detailed mammalian stem cells characterization, its unique life cycle**—** where all tissues and differentiated cells, except for stem cells, are eliminated during weekly regeneration**—** makes stem cell aging the primary driver of observed organismal aging phenotypes. This makes *B. schlosseri* an exceptional model for aging research, a capacity further enhanced in our laboratory by the long-term cultivation and documentation of colonies from their initiation, at the moment of larval settlement over extended time. Some of these colonies have been maintained for over 24 years, having progressed through more than 1,000 asexual life cycles. These characteristics establish *B. schlosseri* colonies as the chordate model with the most documented asexual life cycles in the scientific literature, offering an unparalleled system for aging research. Consequently, in a system where aging is so fundamentally linked to stem cell aging, any reversal of the aging process by interventions like PEC would necessarily involve changes within the stem cells themselves.

Our data show that PEC stimulation improves overall health and enhances stem-cell-mediated budding and regeneration in both young and old *B. schlosseri* colonies, effects reproducible over 4 years of study. The persistence of rejuvenation through numerous asexual cycles in aged colonies strongly implicates alterations within their stem cell populations, as these are solely responsible for the budding process and are the only cells to endure across generations. While our flow cytometry analyses suggest PEC-induced changes in specific cell populations, the current lack of *Botryllus*-specific antibodies and the inherent rarity of stem cells in general circulation limit the resolution of these assays for directly quantifying the stem cells themselves. Therefore, flow cytometry primarily allows us to detect changes in larger, downstream cell populations rather than making definitive assertions about direct PEC effects on the stem cells.

Both transcriptomic and flow cytometric analyses indicate PEC-induced changes in immune cell populations. While myeloid cell populations in *B. schlosseri* are not as comprehensively characterized as in humans, groups of cell populations exhibiting myeloid transcriptomic signatures and immune activity have been identified^32^. Notably, macrophages and their equivalent of myeloid populations are known to reside in a cell population that expands following PEC treatment, paralleling observations where certain myeloid cell populations increase in humans after exercise. These observed changes in macrophage or myeloid-like populations are consistent with the broader organismal rejuvenation observed.

Given that macrophages show age-related functional changes^38^, it is of particular interest that short-term PEC stimulation (3 x 5 min at 150 ppm) in *B. schlosseri* induces a rapid upregulation of a gene signature associated with an M1 macrophage inflammatory cytokine phenotype within 2 hours. This response closely resembles the M1 activation observed in humans and mice following exercise, as defined by the expression profiles of a selected set of genes associated with reactive (M1) and tolerant (M2) phenotypes^16,17^.

Conversely, >24 hours after either exercise (in mammals) or PEC treatment (in *Botryllus*), the transcriptomic signature switches to a tissue-reparative (M2) phenotype, balancing the initial inflammatory response. While the precise mechanisms underlying the similarity between a brief PEC stimulation and the effects of sustained exercise on macrophage activation remain to be fully elucidated, the significant increase in heart rate observed in PEC-stimulated animals may play a role. Thus, despite being distinct interventions, exercise and PEC appear to promote health through similar mechanisms, including the induction of a dynamic shift in macrophage activation.

PEC may induce rejuvenation by resetting cells to a more primitive state. The observed change in *Tgs1* gene expression, a known regulator of telomerase inactivation, is particularly intriguing. Since telomerase activity declines with age and during ovarian maturation^39^, telomerase re-activation has been found to reverse tissue degeneration in aged mice^40^. This mechanism could play a significant role in the PEC-induced rejuvenation we observed.

While PEC induces many of its rejuvenating effects in both young and old *B. schlosseri* organisms, its impact on fertility required nuanced consideration. We observed a definitive increase in gonad production in younger PEC-treated animals. However, a similarly thorough evaluation in older animals (14-24 years) was constrained by the smaller number of these exceptionally aged colonies available for experimentation. There may be a threshold age beyond which the PEC effect on gonad development will not be seen, though we have yet to do a complete series of experiments with large numbers to clarify that age. Further evaluation is necessary to determine if we increase gonad numbers in older populations of animals.

Marine organisms face escalating threats from warming and acidifying oceans, which disrupt critical ecosystems like coral reefs and kelp forests^41,42,43,44^. Warmer temperatures can exacerbate pathogen growth, weaken immunity, and alter host-pathogen interactions, thereby increasing disease prevalence ^45,46,47,48,49,50,51^. Developing methods to bolster marine organism resilience is thus critical. Tunicates are evolutionary survivors, that have adapted to diverse environmental changes over at least 550 million years^34^, further validating their suitability for studying mechanisms that confer environmental resistance. Our observation of PEC-induced rejuvenation in *B. schlosseri*, including indications of enhanced immune function and resistance to aging-related declines, suggests the potential of PEC to fortify organisms against such environmental challenges. Indeed, our PEC data aligns with decades of fieldwork demonstrating the potential of electrical stimulation to promote coral reef regeneration ^31,30^. Electrical structures designed for low-voltage current delivery have shown promise in increasing coral settlement, growth, and survival, including increased resilience to thermal stress^52^. Integrating insights from these extensive field studies with our mechanistic findings in the *B. schlosseri* PEC model will allow for a more rigorous investigation of these protective effects, paving the way for impactful research and potential environmental applications.

The observation of significant and long-lasting rejuvenating effects from a brief PEC session (completed within an hour) holds significant implications for both field applications and potential clinical translation. The minimal time and resources necessary to achieve the desired effect are noteworthy. Therapeutic or environmental benefits could potentially be achieved with minimal cumulative current, thereby reducing demand for battery power or other energy sources. For potential applications in mammalian systems, this could translate to minimal surgical or anesthetic exposure. Existing small, remotely controlled battery-powered devices may prove sufficient for delivering such effective, short-duration treatments.

## Conclusion

We have demonstrated that short-term exposure to low levels of pulsatile electrical current dramatically affects the colonial tunicate *Botryllus schlosseri*. Multiple aspects of the aging phenotype, including zooid size, bud/zooid number, colony survival, and gonad formation exhibit rejuvenation towards a younger state following PEC treatment. Transcriptome analysis suggests contributing pathways, including potential telomerase reactivation and a strong macrophage response that strikingly mimics the exercise-induced macrophage activation observed in mammals. Intriguingly, both exercise and PEC, despite being distinct interventions, appear to utilize similar mechanisms to achieve positive effects on health and survival. These findings and the observed parallels to mammalian responses further demonstrate the utility of *B. schlosseri* as a valuable model for investigating fundamental, evolutionarily conserved biological processes applicable to a wide range of organisms.

## Acknowledgements

Medtronic graciously supplied materials for pacing. This work was supported by a Big Ideas for Oceans grant from the Stanford Oceans Department and Stanford Woods Institute for the Environment to A.V.; Chan Zuckerberg San Francisco Biohub, and a National Institute on Aging grant 5RO1 AG076908 to AV. and ILW; Wu Tsai Human Performance Alliance (WTHPA) to YV and the Gruss Lipper Postdoctoral Fellowship to TL. We thank Stuart Thompson and Steve Palumbi for the helpful discussions.

## Author contributions

Conceptualization: Developing the original idea and research questions. J.D. A.V. E.D. K.G. Y,V. D. S. Methodology: Designing the experiments, developing protocols: J.D. A.V. K.G. E.D. T.L. T.R. K.I. K.P. A.D. N.N. Investigation: Conducting experiments, collecting data. J.D. K.G. T.L. E.D. C.A. K.I. K.P. Software: Developing or modifying software used in the research. Y.V. D.S. T.R. Analysis: Statistical analysis, data interpretation. J.D. A.V. E.D. K.G. T.L. Y.V. Validation: Verifying the reproducibility of results. J.D. A.V. K.I. K.P. T.L. Resources: Providing materials, equipment, or facilities. A.V. I.W. N.N. A.D. Data Curation: Managing and archiving the data. J.D. A.V. Y.V. K.I. K.P. Literature Review: J.D. A.V. E.D. K.G. Writing – Original Draft: Writing the initial manuscript draft. J.D. K.G. A.V. E.D Writing – Review & Editing: Reviewing and editing the manuscript. J.D. K.G. A.V. E.D. T.L. Y.V. K.I. K.P. T.R. N.N. A.D. D.S. I.L.W Visualization: Creating figures and tables. J.D. A.V. Y.V. K.P. K.I. Supervision: Overseeing the research project. J.D. A.V. Funding Acquisition: Obtaining grants or funding for the research. A.V. I.L.W. J.D. K.G.

## Data Availability

RNA-seq data have been deposited at the Sequence Read Archive (SRA) under BioProject PRJNA1219309.

The BioProject and associated SRA metadata are only set to be released on publication, a read-only version is available for reviewers at https://urldefense.com/v3/ https://dataview.ncbi.nlm.nih.gov/object/PRJNA1219309?reviewer=sooash811c8p72trje4h1qr08r;!!Mih3wA!AqXhKUf51mMTugZ6FxE8RICrNdZemYsO1YtnqeiD_kLH3d38PLTsfkurW55Hk01HP9jT9h9WwTfMG7I38uhtZKbi$

The human and mouse exercise data are publicly available with Gene Expression Omnibus ids: GSE104079, GSE27536, GSE120862, GSE83578, GSE111555, GSE99963, GSE144304, and GSE298998.

## Materials and Correspondence

Ayelet Voskoboynik Jos Domen Debashis Sahoo

## Online Methods

### Animal culture and preparation

Mariculture procedures have previously been described^23^. Briefly, *B. schlosseri* colonies were collected from the marina in Monterey, California. Individual colonies were tied to 3 × 5 cm glass slides and placed 5 cm away from and opposite another glass slide in a slide rack. The slide rack was placed into an aquarium, and within a few days, the sexually reproduced tadpoles hatched, swam to the settlement slide, metamorphosed into an oozoid, and matured into the adult body plan. Single colonies are then transferred to individual slides and grown at 18-20°C and under 14h/10h light/dark regimen (6 a.m. - 8 p.m. light/8 p.m. - 6 a.m. dark). Colonies were fed daily using a marine invertebrate diet prepared in the lab as described in^23^. Well-expanded colonies were split into 2 or more subclones and transferred to 2 or more slides, allowing the separate culture of genetically identical animals. Animals were taken for electrical stimulation in stage A and harvested for analysis in stage A or early B. Young colonies used in our experiments ranged from 2 to 18 months. Old colonies ranged from 14 to 24 years in culture.

### ES cell differentiation

E14 ES cells were cultured using standard conditions (DMEM with 10% FCS (GibcoBRL ES cell qualified), 1000U/ml LIF (Chemikon), and fibroblast feeder cells from mouse embryos)^53^. To obtain contracting cardiomyocytes, the cells were plated without feeders or LIF and allowed to grow for one to two weeks until beating cardiomyocyte patches were visible.

### Pulsatile Electrical Stimulus

A Medtronic 5375 pulse generator (Medtronic, MN) was used to PEC-stimulate *Botryllus* colonies. Settings used are as indicated for individual experiments. Animals were routinely stimulated at 20 milliamps (ma) and 150 pulses per minute (ppm) for 5 minutes. This stimulation regimen was repeated three times at 15-20 minute intervals. For some initial experiments, 20 mA, 150 ppm for 5 minutes was applied 6 times with 48-hour intervals. Conditions of the stimulation setup are shown in Supplemental Fig.1a. Briefly, leads with only the last 5mm exposed were positioned on either side of colonies cultivated on glass and submerged in shallow autoclaved seawater sourced from Monterey Bay. Following stimulation, the animals were returned to their mariculture tanks.

Multi-organ duplication was seen in one 5-month-old *Botryllus* colony after extended PEC stimulation. This colony was PEC-stimulated at 150 bpm and 20 mA for 5 minutes every 48 hrs for 12 days. Sixty-nine days later, we saw a doubling of siphons (Supplemental Fig.2) and brains. The doubling of organs was seen in only one asexual generation, unlike what is seen with previously reported bioelectricity-induced changes^1^, which show changes that propagate from generation to generation. While it is unclear if this abnormality was related to the PEC treatment, we limited PEC treatments to 150 bpm and 20 mA for 5 minutes at 20-minute intervals X 3 or 20 mA, 150 ppm for 5 minutes at 48-hour intervals X 6, for the remainder of the experiments.

### Identification of *B. schlosseri* aging phenotypes

Morphological changes such as zooid and bud numbers and the presence or absence of gonads were recorded every 2-5 days during the initial phases following stimulation and monthly afterwards. Pictures and videos of young and old colonies were taken using a Zeiss V12 microscope/Canon camera or iPhone with an in-house produced phone holder and smartphone time lapse controller (Kiss-system TO https://github.com/TomRolander/KISS_Smartphone_Time-Lapse_Controller). Morphological changes were recorded (e.g., pigmentation level, blood vessel shape). Zooid sizes and ampullae length/width ratios were measured using Image J (mm^2^). Heartbeat frequency was determined by analyzing videos of treated and control colonies, focusing on zooid heartbeats. A T-test was used to find significant changes. Graphs were prepared using Graphpad Prism 4.0. Figures were prepared using Pixelmator Pro 3.6.14.

### FACS analysis

Colonies were dissociated with a fine blade into cell suspensions for cell isolation. Cells were filtered through a 70-μm mesh followed by a 35-μm mesh using a sterile 1-ml syringe pump, washed, and collected in staining medium: M199 with 3.3× HBSS. No enzymatic dissociation was used. Cells were resuspended in 1 ml of the staining media, and 1 μl of Alkaline Phosphatase Live Stain (ThermoFisher) was added. Cells were incubated for 30 minutes on ice under dark conditions, and 1 μl of Propidium Iodide (1µg/ml) was added before flow analysis. After gating on propidium iodide (PI) negative cells (using 2D plots owing to the natural fluorescence of *B. schlosseri* cells), cells were analyzed using forward (FSC) and back-scatter (BSC) and using fluorescence from the Alkaline Phosphatase staining on a log scale using a Sony MA900. Flow cytometry data were analyzed using Flowjo.

### Sample-collection for sequencing

Tissue samples were collected from *Botryllus schlosseri* colonies raised in the Hopkins Marine Station Mariculture Facility. They were isolated without food for 20 hours prior to dissection. Colonies were frozen in liquid nitrogen and held at -80°C prior to library preparation. Colonies were sampled from systems at stage A-B.

### Library preparation

RNA was prepared from frozen samples using Zymo Research Quick RNA MIcro Prep Kit # R1050 and cleaned using Zymo Research RNA Clean and Concentrator #R1015. Samples were analyzed on an Agilent QC 2100 Bioanalyzer to determine quality prior to library preparation. cDNA was prepared using the Nugen Ovation RNA Sequencing System V2, #7102, and cleaned using the QIAGEN QIAquick PCR purification kit, #28104, as recommended in the protocol and analyzed on the Agilent QC Bioanalyzer. If needed, samples were size-selected using Zymo Research Select-A-Size DNA Clean and Concentrator #D4080 prior to barcoding. The final library was prepared using NEB NEBNext Ultra II DNA Library Prep Kit #27645 and barcoded using NEBNext Multiplex Oligos for Illumina #E6609S. All magnetic bead purification was accomplished using BullDogBio CleanNGS RNA and DNA Spri Beads #CNGS005. Samples were then analyzed using the Agilent QC 2100 Bioanalyzer to determine the concentration of each sample prior to sequencing. On average, 12 million 2×150 bp reads (Illumina Nextseq 2000 P3) were sequenced for each library.

### Transcriptome analysis

To study molecular changes associated with rejuvenation, we pulsed young (373 to 465-day-olds) and old (7,668-day-olds) colonies, generating comprehensive whole-transcriptome sequence data from 47 samples. Both PEC-treated and control colonies were sampled. Young colonies were sampled at 5 time points post-treatment: 2 hours, 24 hours, 13, 49, and 132 days. Old colonies were sampled at 3 time points: 24 hours, 41 days, and 93 days. All specimens were sampled at the same developmental stage, stage A. Bulk RNA-seq libraries were prepared and sequenced on an Illumina NextSeq 2000 P3 platform. Reads were aligned to the *Botryllus* genome, and gene expression was measured as described in ^23^. Reads were trimmed (trim galore) and aligned to the *Botryllus* genome^12^ using STAR^54^. Gene count tables were created using HTseq. Samples with less than 10,000 total counts were removed. DEseq was used to identify differentially expressed genes with FDR<=0.05 and a Log fold change cutoff absolute value >2. Heatmaps were created based on the Log fold change values of selected genes.

### Gene orthology

Gene orthology is based on sequence similarities between the *B. schlosseri* gene models and human and mouse gene annotations (BLAST score smaller than e-10), as described in the *B. schlosseri* genome paper^12^. All discussions regarding genes are based on sequence similarities alone.

### Macrophage Polarization Scores

The M1/M2 macrophage polarization state analysis was done as previously described^17^. In short, a previously identified macrophage gene set^16^ of 338 genes, which accurately separated M1 vs. M2 macrophages in both training and validation-independent clinically prescribed macrophage datasets (M1 = 48 genes, M2 = 290 genes), was used initially. Of those 338 genes, 112 had a homologous *Botryllus* gene. When multiple *Botryllus* Gis were homologous to the same gene, all such Gis were used, leading to 13 M1 Gis and 133 M2 Gis in the *Botryllus* analysis (SI Table 2). A combined score was calculated from these genes. The genes within each cluster were initially normalized and then averaged to derive the score. The normalization of gene expression values followed a modified Z-score approach centered around the StepMiner threshold (formula = (expression – (SThr + 0.5))/3∗stdev)^55^. A weighted linear combination of the cluster averages was employed to generate a score for each sample. Subsequently, the samples were organized based on this linearly combined score. The direction of the pathway was determined by the connection from an M1 cluster to an M2 cluster, with M1-associated genes being given a negative weight and M2-associated genes a positive weight. A noise margin for this composite score was computed, adhering to the same linearly weighted combined score with a 2-fold change allowance (±0.5 around the StepMiner threshold). In Fig. 5a, the scores for GSE27536 were calculated with only the M1-associated genes due to the disease status of all the samples in those datasets. For the *Botryllus* control vs the 2Hr-post-PEC treatment analysis, only the M1-associated genes were used to calculate the scores, as the PEC stimulus was also considered stress-inducing in the samples. The long-term (control vs 300-1000 Hrs and 1000+ Hrs) *Botryllus* analysis used both the M1 and M2 associated genes.

## SUPPLEMENTAL DATA

### Supplementary Figures

**Supplemental Information Fig 1.**
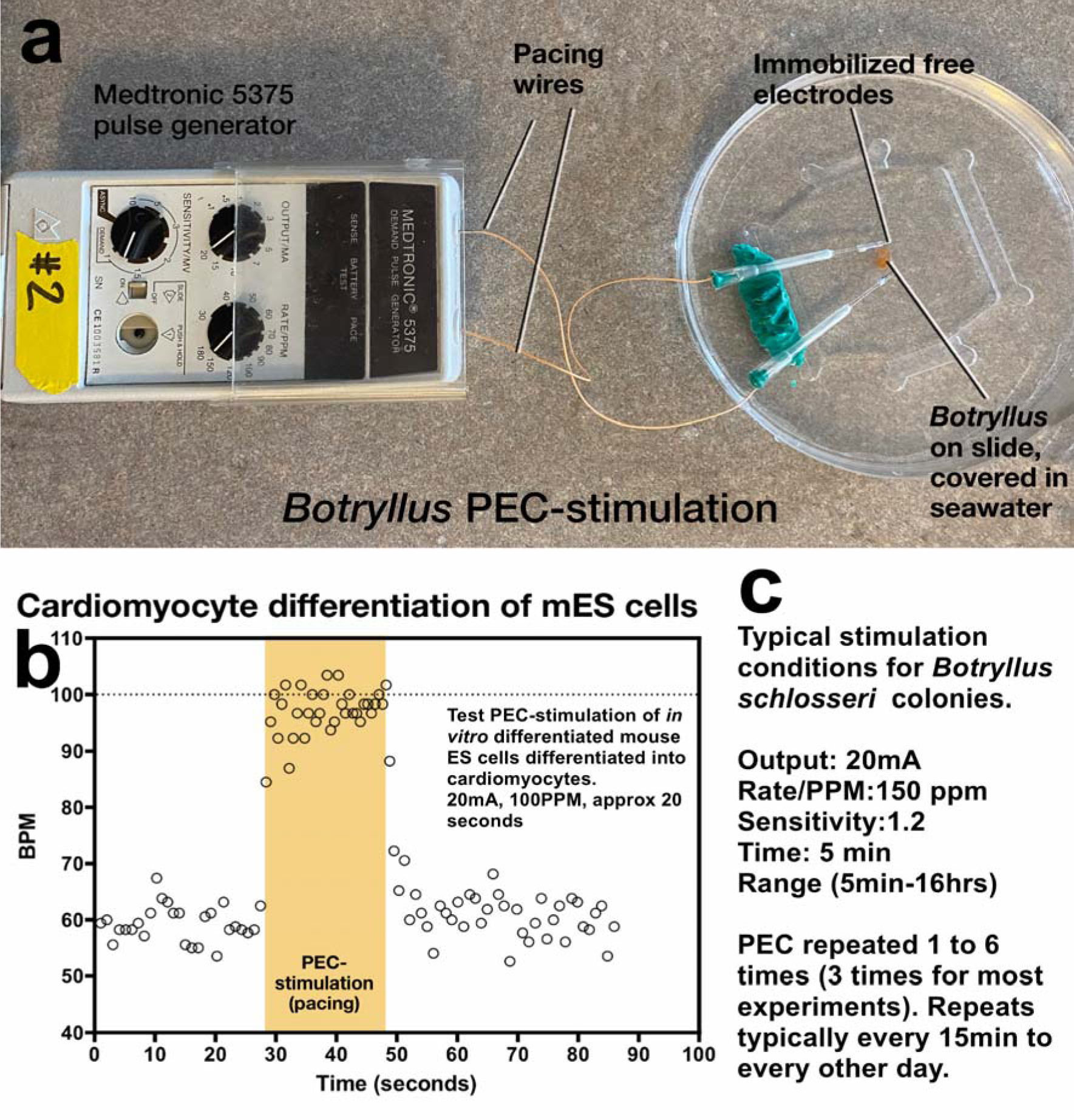
Pulsatile Electrical Stimulation set up for *Botryllus schlosseri*. (a) Experimental setup. *Botryllus* colonies growing on glass slides facing up are submerged in sterile seawater. Pacing wires are positioned on either side of the colony, with only the distal 5 mm of the wires exposed. Following PEC treatment, colonies are returned to their regular mariculture tanks. (b) Validation of pulse generator output. A test run using Medtronic 5375 pulse generator confirms that the contraction frequency of cardiomyocyte patches adapts to the frequency set on the Medtronic device. (c) Typical PEC treatment parameters used for *Botryllus* colonies stimulations

**Supplemental Information Fig 2.**
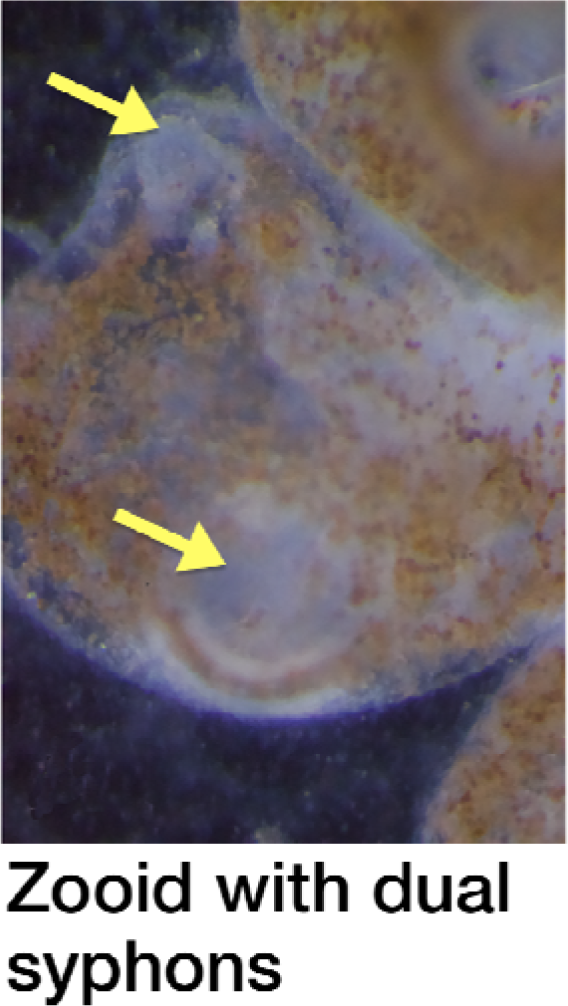
Multi-organ duplication in *Botryllus schlosseri* following extended PEC Stimulation. This figure shows a 5-month-old *Botryllus schlosseri* colony exhibiting multi-organ duplication after extended PEC stimulation. The colony was treated with PEC at 150 bpm and 20 mA for 5 minutes every 48 hours for a total of 12 days. Sixty-nine days after the treatment concluded, the colony displayed a doubling of siphons and brains.

Supplemental Information Video 1 | **Video of blood circulation in a young control colony and PEC treated colony (18 days post treatment).** This video compares blood circulation in a young control colony and a PEC-treated colony 18 days after treatment. Both colonies exhibit vigorous blood flow and heart function, with even greater activity in the PEC-treated colony (see also Fig 2a for quantification). Note the different magnifications: 33.5x for the control colony 6964a and 83× for the treated colony 6964d. Healthy young colonies, such as those shown here, do not exhibit the same dramatic changes in blood flow observed in older colonies (see Video 2)

Supplemental Information Video 2 | **Video of blood circulation in a 21 years old colony before and after PEC treatment.** This video shows a 21-year-old colony before and after PEC treatment, demonstrating the change in extracorporeal vasculature, ampullae density, and blood flow. Note the narrow vessels, densely filled ampullae and sluggish blood flow in the colony before treatment, compared to the wider vessels, less densely filled ampullae, and more robust blood flow 225 days after PEC treatment.

**Supplemental Information Table 1.**
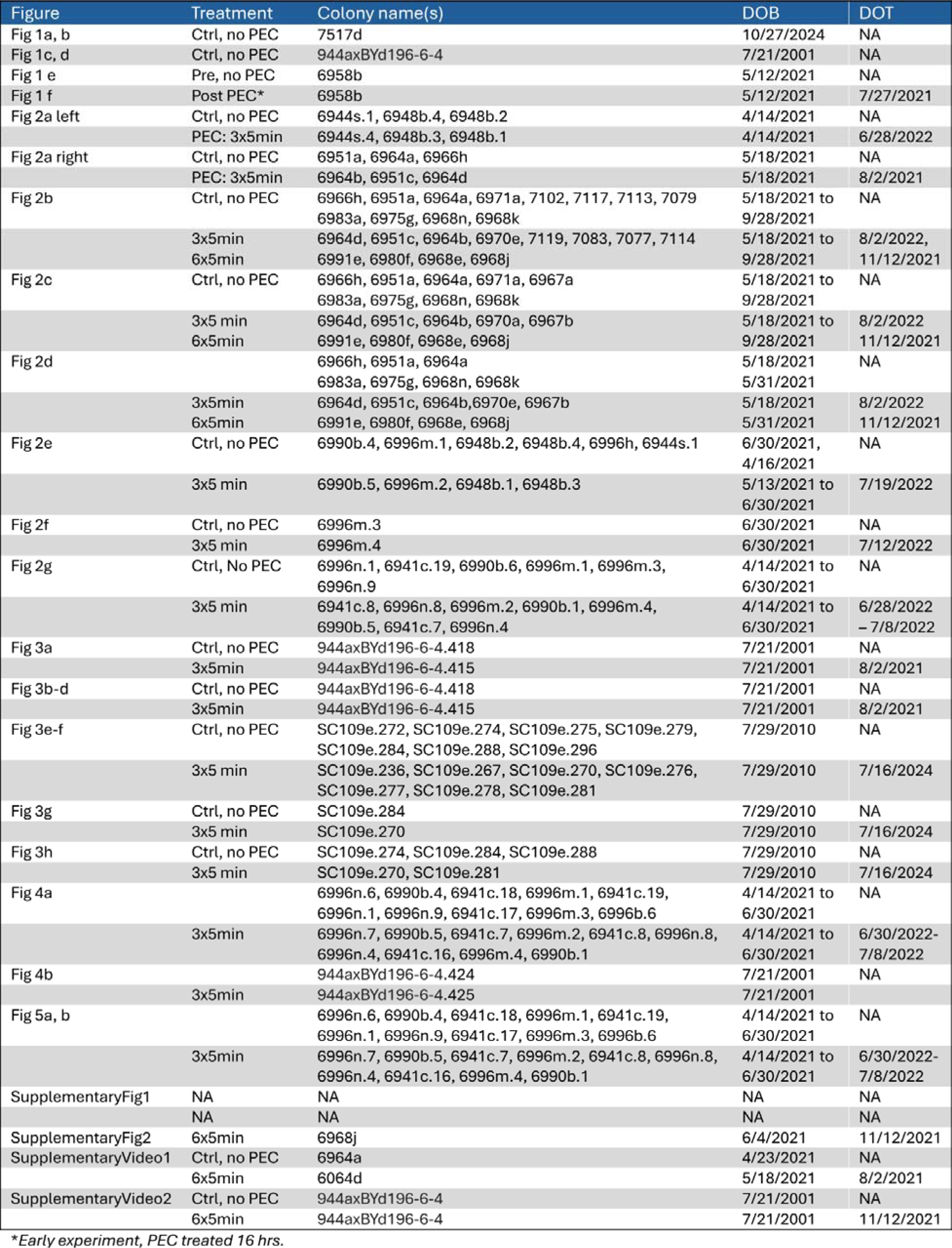

### Identification of Organisms Used in Figures

This table provides information on the organisms used in the figures, including colony name (identity number), date of birth (DOB), date of PEC treatment (DOT), and treatment details.

**Supplemental Information Table 2.**

### Data and Analyses

This series of Excel spreadsheets contains the data and analyses related to the RNA-Seq.

▭ **Sequenced Samples Metadata**: Information about the sequenced samples, including experimental group, treatment, age, and other relevant metadata.
▭ **DESeq Group Annotation**: Annotation file for DESeq analysis, specifying the experimental design and comparisons.
▭ **RNA-Seq Counts Table**: Raw read counts for each gene in each sample.
▭ **Group 1 DESeq Results**: Differential gene expression results for Group 1 comparison.
▭ **Group 2 DESeq Results:** Differential gene expression results for Group 2 comparison.
▭ **Group 3 DESeq Results**: Differential gene expression results for Group 3 comparison.
▭ **Group 4 DESeq Results**: Differential gene expression results for Group 4 comparison.
▭ **Group 5 DESeq Results**: Differential gene expression results for Group 5 comparison.
▭ **Group 6 DESeq Results**: Differential gene expression results for Group 6 comparison.
▭ **M1 and M2 Botryllus Gene Set**: List of genes associated with M1 and M2 macrophage polarization in Botryllus.

